# Blood-based transcriptomic signature panel identification for cancer diagnosis: Benchmarking of feature extraction methods

**DOI:** 10.1101/2022.03.13.483368

**Authors:** Abhishek Vijayan, Shadma Fatima, Arcot Sowmya, Fatemeh Vafaee

## Abstract

Liquid biopsy has shown promise for cancer diagnosis due to its minimally invasive nature and the potential for novel biomarker discovery. However, the low concentration of relevant blood-based biosources and the heterogeneity of samples (i.e. the variability of relative abundance of molecules identified), pose major challenges to biomarker discovery. Moreover, the number of molecular measurements or features (e.g., transcript read counts) per sample could be in the order of several thousand, whereas the number of samples is often substantially lower, leading to the curse of dimensionality. These challenges, among others, elucidate the importance of a robust biomarker panel identification or feature extraction step wherein relevant molecular measurements are identified prior to classification for cancer detection. In this work, we performed a benchmarking study on eleven feature extraction methods using transcriptomic profiles derived from different blood-based biosources. The methods were assessed both in terms of their predictive performance and the robustness of the biomarker panels in diagnosing cancer or stratifying cancer subtypes. While performing the comparison, the feature extraction methods are categorised into feature subset selection methods and transformation methods. A transformation feature extraction method, namely PLS-DA, was found to perform consistently superior in terms of classification performance, while a feature subset selection method, namely Ranger, selected feature sets that were the most robust to sub-sampling in terms of consistently selecting the same features. As part of the benchmarking study, a generic pipeline has been created and made available as an R package to ensure reproducibility of the results and allow for easy extension of this study to other datasets.

## Introduction

The traditional biopsy method for identifying cancer requires a surgical procedure to obtain a tissue sample from the patient. As an alternative, the recently popularised liquid biopsy method involves the analysis of non-solid tissue such as blood, saliva and urine to identify molecules originating from the tumour [1]. A wide variety of analytes such as circulating tumour cells (CTCs), circulating nucleic acids including circulating tumour DNA (ctDNA), tumour-derived fraction of cell-free DNA (cfDNA) in the plasma, as well as cell-free RNAs (mRNAs, long non-coding RNAs and microRNAs), extracellular vesicles, tumour-educated platelets, proteins and metabolites are present in a range of bodily fluids [2]. These have the advantage of being easy to obtain, minimally invasive and cost effective, and allowing for longitudinal assessment. In addition, studies have shown that they are also capable of capturing the clonal heterogeneity of the disease over time [3]. Blood-based biomarkers thereby overcome the limitations of tissue acquisition and stand out as the holy grail for cancer diagnosis and monitoring [4].

One of the analytes that has attracted enhanced attention as a source of biomarkers are Extracellular Vesicles (EVs). EVs are secreted from many cell types and play important roles in intercellular communications. EVs carry a range of biomolecules that reflect the identity and molecular state of their parental cell and are found in biological fluids. High-throughput molecular profiling studies (i.e., omics) have focused on characterisation of the molecular cargo (e.g., RNA transcripts) of EVs [5]. EVs are increasingly being utilised in disease diagnosis as they are considered to carry valuable information about the disease state [5]. Beyond EVs, tumour educated platelets (TEPs) or platelets that have been ‘educated’ through the transfer of tumour specific RNA, have recently been discovered as blood-based biosources that hold promise in cancer diagnosis. Together with computational modelling, the RNA profiles of TEPs have been used to distinguish between multiple cancer types with over 70% accuracy [4, 6].

However, liquid biopsy comes with its own set of challenges which limits its usage as a standardized clinical tool. The concentration of tumour originated molecules in blood and other body secretions is very low, which makes their extraction a tedious process and adds substantial noise to the resultant data. There also exists the issue of variability across different samples due to the method used for blood collection and choice of sequencing technology such as RNA-seq or microarray [2]. Moreover, the composition of the biosources are heterogeneous across samples derived from different patients, adding another level of variability to the data generated. Another major issue is that the number of transcripts is roughly in the thousands, whereas the number of samples are often less than a hundred.

Computational cancer diagnosis from high-throughput omics data usually involves two steps: 1.) identification of a biomarker panel, i.e. a set of transcripts which are good indicators of a condition 2.) classification of patients into conditions of interest (e.g. cancer vs normal) based on the biomarker panel. The first step involves extracting a set of relevant identifiers or features contributing to a condition, which is called the *feature extraction* step. Specifically for liquid biopsy based cancer-diagnosis with small sample size and high-dimensional feature space along with the inherent variability in features (e.g., transcript expression measures) among patients, a robust feature extraction procedure is necessary prior to the cancer classification task. Feature extraction can reduce the dimensionality and sparsity of data and enhance model generalisation by reducing the chance of overfitting the model. Therefore, careful consideration should be given to the feature extraction method in order to improve the prediction performance while maintaining robustness across different patient cohorts.

The main objective of this study is to compare feature extraction methods for cancer diagnosis from blood-based liquid biopsies. The comparison of feature extraction methods is done both in terms of their predictive performance and robustness. A feature extraction method is considered to be robust if the performance of the prediction task after feature extraction is similar across multiple sub-samples of the patients. This has been achieved both by directly comparing the performance, and also by verifying if the feature extraction method selects similar features across different randomly sub-sampled training cohorts. Due to sample heterogeneity, it is likely that the transcripts found to be predictive in one cohort, only partially exist in another cohort of patients with the same disease, thereby making the biomarker panel irreproducible and not robust. Therefore, using the full transcriptomic profile after applying some transformations may enhance the reproducibility compared to the conventional approach of selecting a small subset of transcripts as the signature panel. Taking this into consideration, this study compares feature subset selection methods that select a subset of relevant transcripts as a biomarker panel, against transformation-based feature extraction methods that use the entire set of transcripts (often after an initial filtration of invariant features, e.g., transcripts with zero counts across most of the samples), and transform the features into a latent or embedding space. Oftentimes, the dimensionality of the latent space is chosen to be lower than that of the original feature space, achieving dimensionality reduction or information compression. The comparison of feature extraction methods is performed by building a generic pipeline to run any transcriptomics datasets on multiple feature extraction methods and classification models.

Previously efforts have been made to systematically compare and evaluate methods for feature selection [7–9] and dimensionality reduction [10] in biomarker discovery from omics data, however there is no study specifically focused on liquid biopsy. As far as is known, this is the first work to systematically evaluate and compare the impact of feature extraction methods on liquid biopsy datasets. Moreover, this work also stands out from the previous ones [7–9] by comparing feature subset selection methods and transformation-based feature extraction methods. Also, the generic pipeline and the associated R package created as part of this study is made publicly available. This will be useful for further benchmarking studies on omics data and for easy extension of the study to other feature extraction methods or new datasets.

## Materials and methods

### Datasets

Assessing the performance of feature extraction methods for cancer prediction has been performed using publicly available transcriptomic datasets for GBM (Glioblastoma, the most aggressive variant of brain tumour), Lung Cancer, Colorectal Cancer, Breast Cancer and Pancreatic Cancer. The details of the datasets used in this study are provided in Table 1. The ‘Dataset Name’ field is the name assigned to refer to each dataset in this study.

**Table 1.**
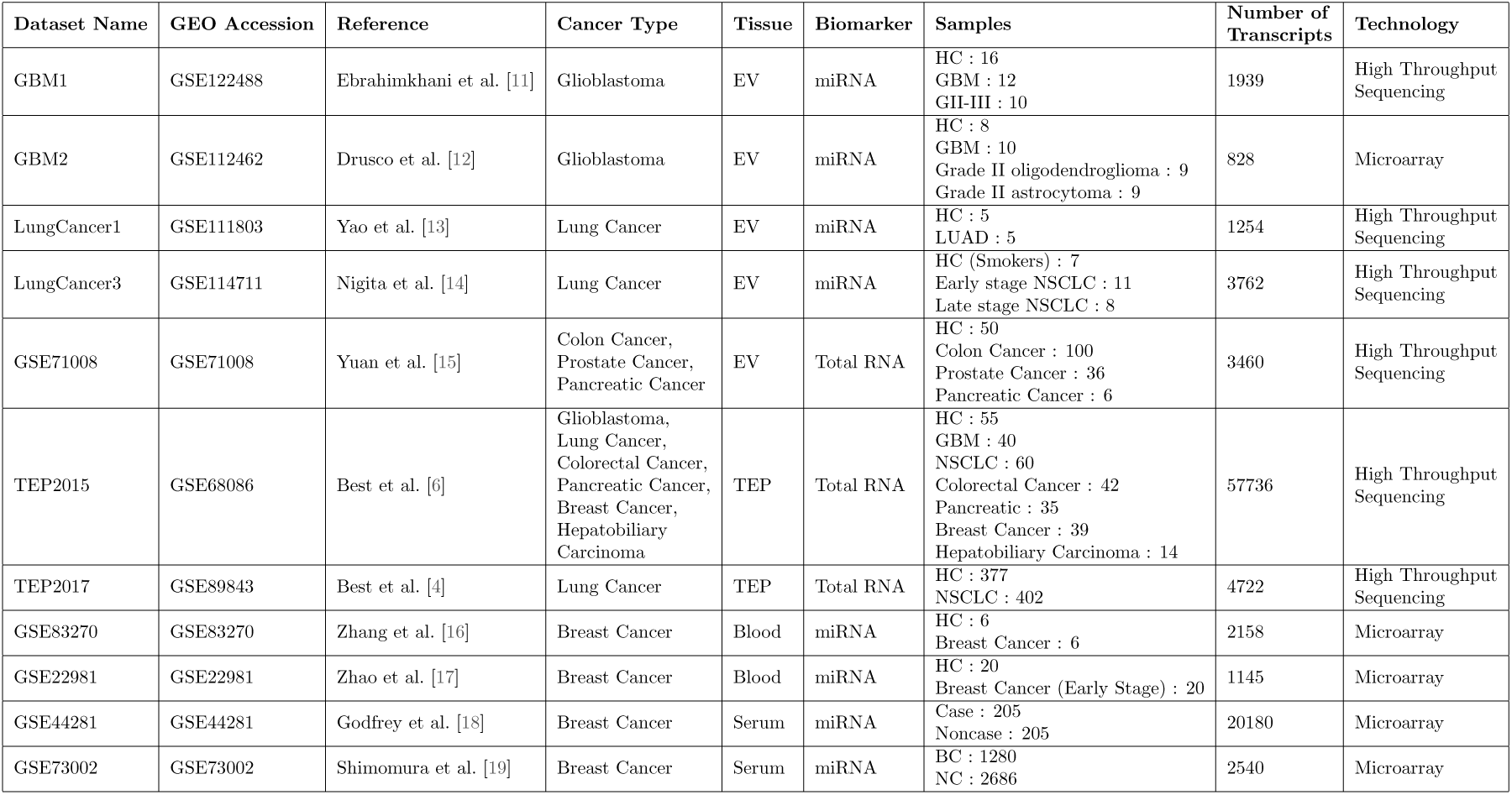
Datasets Used

The datasets were identified primarily using BBCancer [20], which is a web-accessible aggregation of various biomarker datasets. The source datasets were retrieved from the Gene Expression Omnibus (GEO) repository with the accession numbers provided in Table 1. Using the source datasets was necessary because BBCancer combines datasets across sequencing technologies. Moreover BBCancer separates out Healthy Controls (HC) from all the source datasets and provides it as a single combined set. This was not favourable for binary classification of cancer types against healthy controls. Apart from BBCancer source datasets, a few other datasets were directly obtained from our former study or by searching GEO including GBM EV dataset from Ebrahimkhani et al. [11] (GSE122488) and TEP datasets for multiple cancer types from Best et al. [4, 6] (GSE68086, GSE89843). The inclusion criteria for datasets were the availability of at least 10 samples, availability of HC data and the availability of blood-based samples (tumour tissue-based data were excluded). The datasets used in this study are microRNAs (miRNAs) or total RNA profiles from Extracellular Vesicles (EVs), whole blood, serum or Tumour Educated Platelets (TEPs).

### Computational Pipeline

The overall pipeline developed as part of this study is shown in Figure 1. The pipeline was implemented in R language [21], using various R packages. To facilitate obtaining results easily from any transcriptomic datasets in the future, the pipeline has been developed in a generic manner and made available as an R package named ”FEMPipeline”. The code and associated R package can be freely accessed from GitHub : https://github.com/abhivij/bloodbased-pancancer-diagnosis.

**Figure 1.**
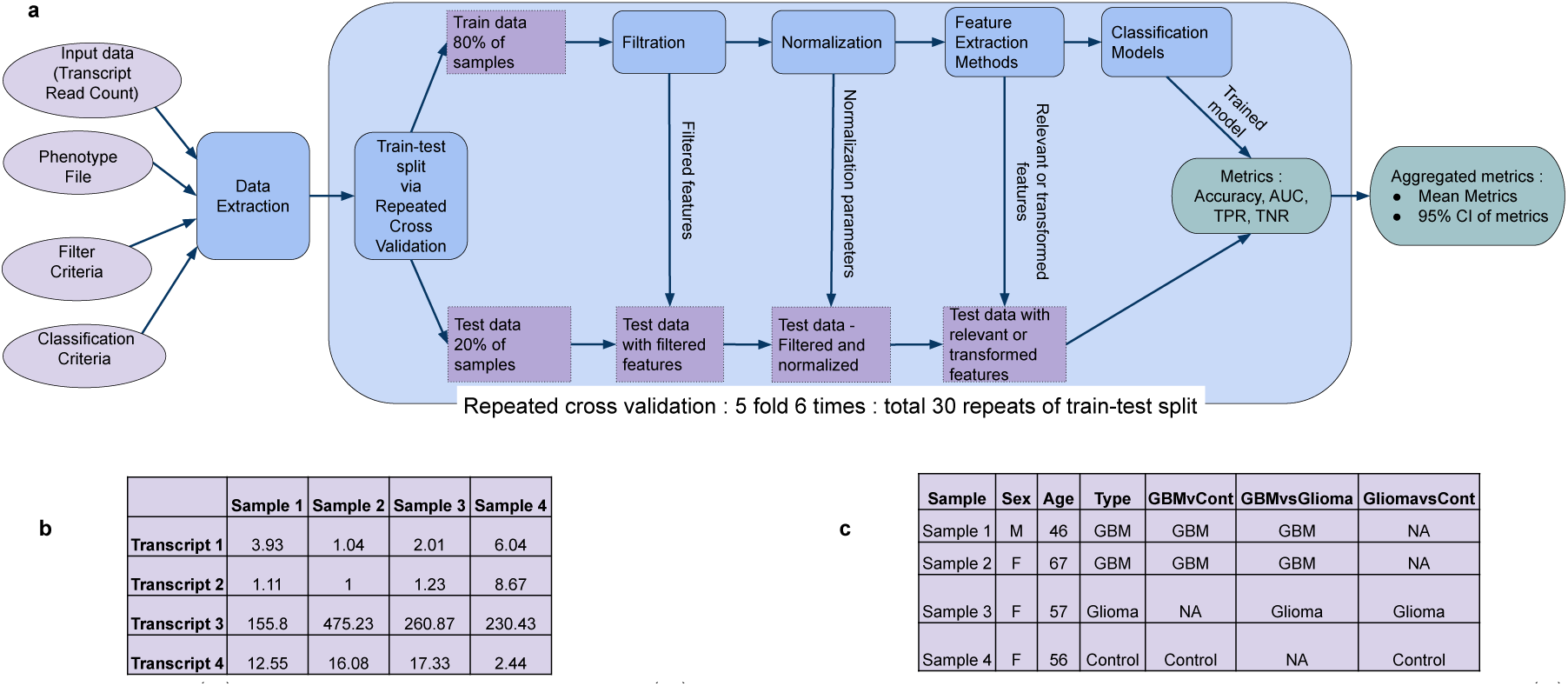
(a) Computational pipeline (b) Schematic example of a transcriptomics input file (c) Schematic example of phenotype file

The following sections describe the different steps in the pipeline.

#### Data Extraction

This step is intended for converting the input data into a common format across all datasets. It extracts only those samples that correspond to the required classes for the binary classification task. Inputs to this step are :

1. Transcriptomics data file : a tab-separated input transcript data that is a matrix where rows represent transcripts and columns represent samples. Each element of the matrix represents read counts (for RNA-sequencing technology) or relative gene expression levels (for microarray technology) as demonstrated in the table in Figure 1 b.
2. Phenotype file : a tab-separated manually created sample specific information.An example of a phenotype file is given in the table in Figure 1 c. This file should contain a row for each sample in the transcriptomics data file, and should also contain columns to be used as filter criteria or classification criteria.
3. Classification criteria : a column in the phenotype file (eg: GBMvsCont), that contains for each of the samples, one of the two classes for the binary classification task (eg: GBM, Control), or NA for samples of other classes.
4. Filter criteria : is an optional input containing an expression for selecting a subset of samples, eg: *expression*(*Age* > 55 & *Sex* == *M*). If filter criteria needs to be specified, then the corresponding variables (Age and Sex as in the example) should be present as columns in the phenotype file. Although filter criteria is supported in the pipeline, it has not been used in this study since the number of samples is quite small.

Outputs from this step include two files :

1. Filtered input data : is a subset of the input transcriptomics data file, containing only the samples that do not have NA for the classification criteria, and those that satisfy the filter criteria.
2. Output labels : is a two column file mapping each sample id in the Filtered input data against a class label. The class label corresponding to each sample is selected from the Classification Criteria column (eg : GBM, Control).

After data extraction, a total of 23 datasets were obtained from the original 11 datasets (Table 1) using multiple classification criteria. These 23 datasets were used as separate datasets for the rest of the pipeline. The 23 datasets are listed in Table 2. In the ’Extracted dataset name’ field, the part before the underscore indicates the original dataset name and the part after it indicates the classification criteria based on which the specific subset of data was extracted.

**Table 2.**
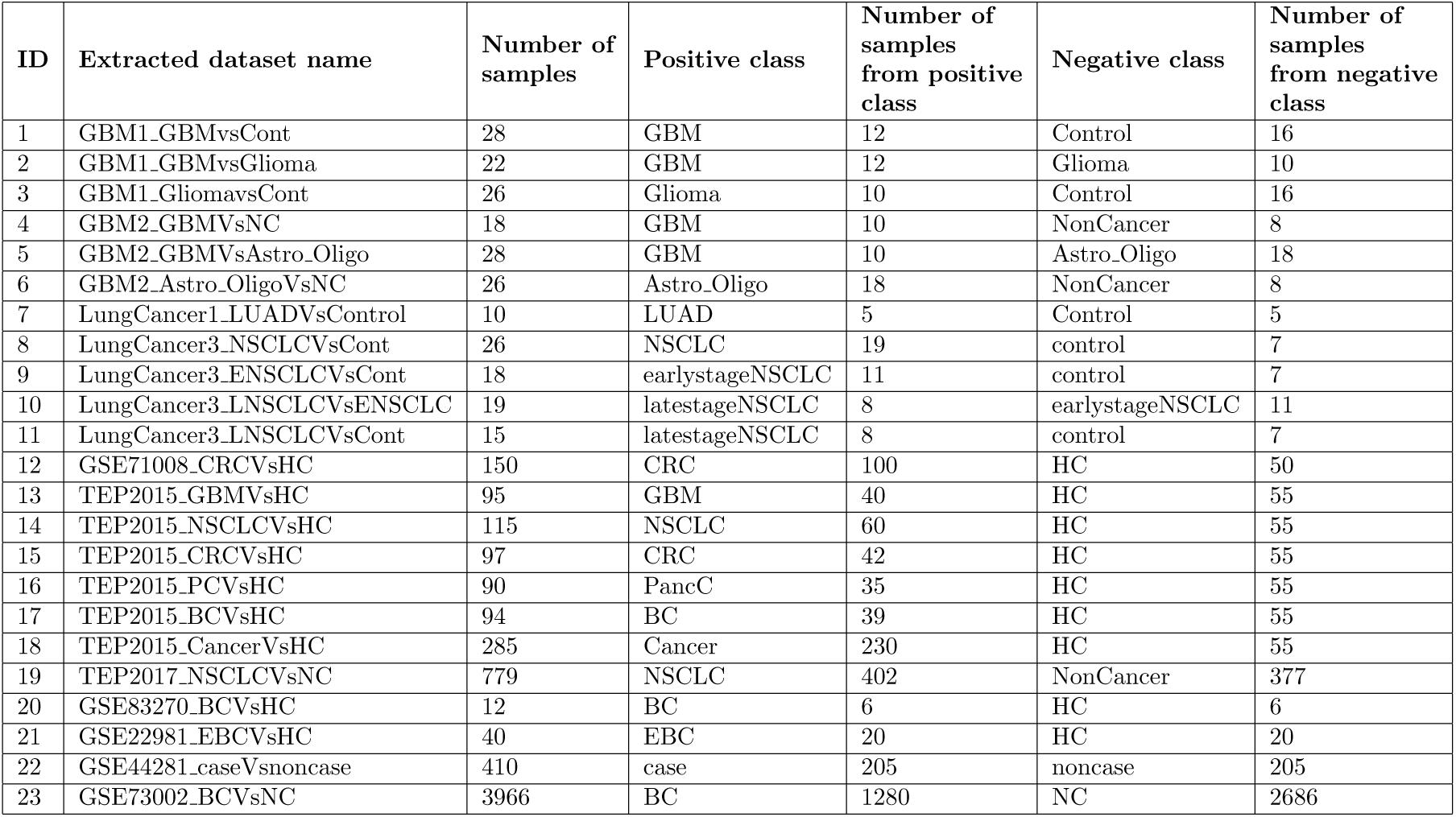
Datasets after Data Extraction

#### Repeated Cross Validation

To reliably estimate the performance of the feature extraction methods and classification models, k-fold cross validation was used to split each dataset into training and test sets. Accordingly, five-fold cross validation was adopted, wherein patients were randomly partitioned into 5 non-overlapping partitions or folds, with one fold reserved as the test set each time and the remaining four folds used as the training set, giving an 80:20 Train-Test split and 5 different training-test combinations. To account for initial random partitioning and to enhance generalisability of estimated model performance, five-fold cross validation was repeated 6 times, providing a total of 30 data-splits (6 repetitions times 5 folds). The remaining steps in the pipeline from preprocessing to classification modelling were repeated for each of these data splits. The parameters in each of the remaining steps in the pipeline are computed using the train data, and applied on the test data to avoid any data leakage or sharing of information between the test and training data-sets.

#### Preprocessing

This step includes filtration and normalisation as detailed below:

**Filtration** is a common preprocessing step for identifying and removing RNAs that present in too small quantities to make any significant differences [4, 11]. In this work, FilterByExpr function from edgeR package [22] in R was used. This tests if the normalised expression values are greater than or equal to a cutoff value (based on a default minimum count and median library size) in atleast a minimum sample size.

**Normalisation** is the process of correcting experimental variations and bias in transcriptomic data to make expression values comparable for valid and reliable downstream analyses [23]. The normalisation method used in this study is Norm Log count-per-million (CPM). This involves computing the log CPM of the transcripts, using edgeR package [22], and then standard scaling (i.e. scaling centered around mean with unit standard deviation) each of the samples. This is followed with standard scaling of each of the transcripts using parameters(mean and standard deviation) computed from the training data, using caret package [24].

#### Feature Extraction

Feature Extraction Methods (FEMs) form the crux of this study. Three strategies of FEMs were used including 1) Using all features, 2) Feature subset selection methods, and 3) Transformation methods, as detailed below:

#### Feature Subset Selection Methods

select a subset of the existing features to be the most relevant ones based on some criteria. These methods often fall into three categories of:

1. Filter Based Methods This subset of methods identifies feature relevance based on intrinsic properties of data by calculating a feature relevance score and removing low-scoring features [25]. These can be further subdivided into univariate and multivariate methods.
  a. Univariate filter based methods are those that consider each feature separately and ignore feature dependencies. Two commonly used methods belonging to this category, namely T-Test and Wilcoxon Rank Sum, were assessed in the pipeline. T-test is a parametric method, i.e., it requires the normality assumption whereas Wilcoxon Rank Sum is a non-parametric test. For both the methods, the relevant features are selected as those that have a p-value less than 0.05 when comparing the samples from the corresponding two classes (e.g., : Cancer vs. Healthy). The stats package in base-R [21] was used for these methods.
  b. Multivariate filter methods overcome the limitations of univariate methods by modelling feature dependencies. Minimum Redundancy Maximum Relevance (MRMR) method belonging to this category was evaluated in this pipeline. This method selects features that are maximally dissimilar to each other so as to reduce redundancy, while being maximally relevant to predicting the phenotype [26]. The number of features to be selected should be prespecified for MRMR. In this study, 30 and 50 features were selected with MRMR. The package mRMRe [27] in R was used.
2. Wrapper Methods In this type of methods, a search procedure among various subsets of features is carried out and the feature subsets chosen in each step is evaluated using a classification algorithm i.e. the search procedure is wrapped around the classification algorithm [25]. The pipeline implemented in this work uses Genetic Algorithm with Random Forest as optimization criteria, which is a wrapper method. Genetic Algorithm (GA) is inspired from the phenomenon of adaptation that occurs in nature, by moving from one population to the next using a kind of natural selection based on a fitness score, together with the genetics inspired operators of crossover, mutation and inversion [28]. For the task of feature selection, individuals of the population are the features themselves, being selected or not selected. The performance of the selected features on Random Forest classification algorithm is used as the fitness score. The caret package [24] was used for this. The default GA optimisation strategy in caret package of using Repeated Cross Validation, took several days for execution even on small datasets using our local high-performance computing platform. Therefore, Cross Validation was instead used as the optimization strategy within GA.
3. Embedded Methods Feature selection is performed as part of a classification algorithm in this variety of method [25]. Two embedded methods were explored in the pipeline.
  a. Random Forest Recursive Feature Elimination: In this method, the importance of each of the features is determined based on model performance using all features. After that, the importance of features are recalculated based on model performance with different subsets of features, and finally the best set of features among all the subsets is reported. Caret package [24] was used. The default number of features in the package are 4, 8 and 16, which are substantially less than the total features in the datasets used. Instead, subsets of size in powers of two from 2 until the total number of features [2, 4, 8, …, power of 2 less than total features] were used.
  b. Features selected by Ranger : Ranger [29] is a fast version of Random Forest designed for high dimensional data. In this method, those features for which Ranger assigns non-zero importance are selected. The Ranger package [30] was used for this method. Feature importance computation was performed using ’impurity corrected’ option. Impurity measure is Gini index for classification and its corrected version is unbiased in terms of number of classes and class frequencies [30, 31].

#### Transformation Methods

These methods make use of the entire set of features after applying some transformations. In this study, the transformation methods explored were dimensionality reduction methods. Each of the following dimensionality reduction methods were explored with embedding of size 2 and 5.

1. PCA: Principal Component Analysis is a linear transformation of input features such that most variance in the data is captured by the first transformed component, second most variance by second component and so on. stats R package in base-R [21] was used.
2. Kernel PCA : Non linear version of PCA. Kernlab package [32] was used for this method.
3. UMAP : General non-linear dimension reduction by modelling the manifold with a fuzzy topological structure [33]. The implementation from umap R package [34] was used. One of the hyperparameters used in UMAP is the number of nearest neighbours (n neighbors) with a default value of 15. In some of the datasets used in this study, the sample size in the training set is smaller than 15, and in those cases n neighbors was set to half the number of samples in train set.
4. PHATE : A recent method that captures both local and global nonlinear structure using information-geometric distance between data points [35]. PhateR package [36] was used for this method.
5. PLS-DA (Partial Least Square Discriminant Analysis) : Dimensionality reduction method that takes class labels into account. This method is considered as a supervised version of PCA. In the method, the transformation is such that the covariance between data and labelling is maximally preserved in the first component [37]. PLS-DA implementation from Caret package [24] was used.

#### Classification Models

The features provided by the feature extraction methods were evaluated through prediction using six commonly used classification models of three different types, namely logistic regression based (Simple Logistic Regression, L1 Regularized Logistic Regression, L2 Regularized Logistic Regression), SVM based (Sigmoid Kernel SVM, Radial Kernel SVM) and decision tree based (Random Forest). Logistic Regression is a classification algorithm that separates classes of interest by fitting a sigmoid (or logistic) function. The regularized version of logistic regression aims to reduce overfitting. Support Vector Machine (SVM) separates the classes of interest by finding a mapping (called kernel) to a higher dimension, and then identifying the best separating hyperplane in that higher dimension. Random Forest is a classification algorithm that separates classes based on the majority vote by large number of decision trees trained on different subsets of data. The R package glmnet [38] was used for Logistic regression models, e1071 package [39] for SVM and randomForest package [40] for Random Forest.

#### Parameter Values

Following the recommendations for benchmarking studies from Weber et al. [41], default parameter values were used for each of the FEMs and classification models to prevent favouring of some methods. Exceptions to this are for genetic algorithm, RF-RFE and UMAP as previously mentioned in the respective method descriptions.

#### Aggregated metrics

For each train-test split, the following metrics were computed on the test data : accuracy, True Positive Rate (TPR), True Negative Rate (TNR), Area Under ROC Curve (AUC). True Positive Rate is the ratio of predicted positive class against all the positive class. True Negative Rate is the ratio of predicted negative class against all the negative class. ROC (Receiver Operating Characteristics) curve is a plot of true positive rate against false positive rate for different values of classifier thresholds [42]. ROCR package [43] was used for AUC and Caret package [24] for TPR and TNR. These metrics were aggregated across the 30 models trained and tested as part of the Repeated Cross Validation. The mean values of each of these metrics were computed to obtain an overall measure. To obtain a sense of the variability in performance, the 95 percent Confidence Interval (95 CI) for the metric values across the 30 iterations was also computed. Since the AUC is independent of a threshold value for the prediction, it is more reliable, therefore the evaluation of results was based solely on Mean AUC and 95 CI AUC for the 30 iterations.

### Computational Resources

The Katana computational cluster, supported by Research Technology Services, UNSW Sydney [44], was used for the execution of the pipeline on the 21 datasets. The job execution in Katana utilized an array job allocating 16 CPUs, 124 GB memory and wall-time ranging from 48 to 200 hours depending on the datasets.

## Results and Discussion

The evaluation of FEM performance was conducted by comparing the Mean AUC scores for each of the classification models on the different datasets.

### Mean AUC

The Mean AUC on each of the 23 datasets for all the feature extraction methods and for each of the 6 classification models are represented as a heatmap in Figure 2.

**Figure 2.**
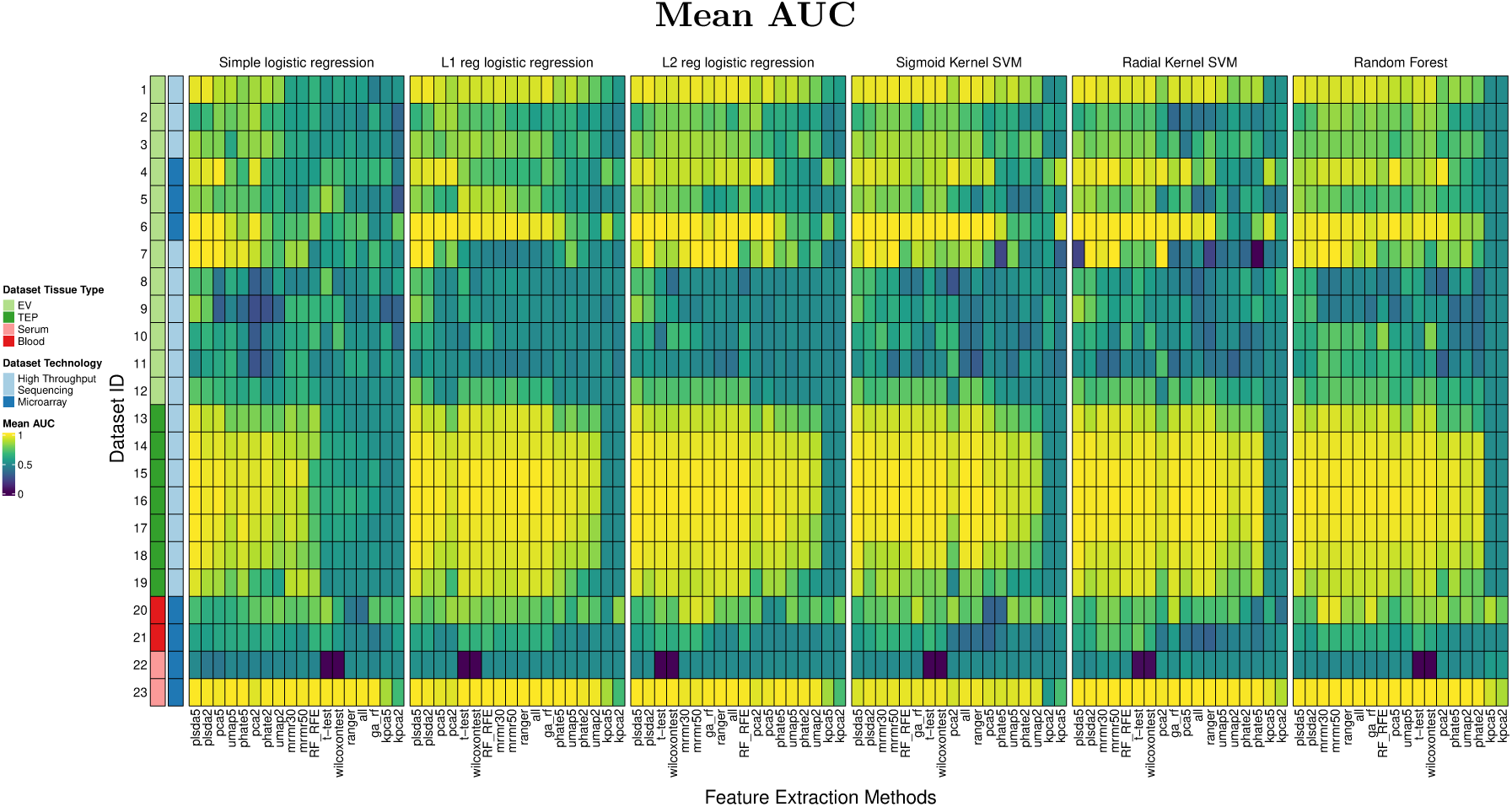
Heatmap of Mean AUC values

For transformation based FEMs, performance with embedding sizes of 2 and 5 are shown. pca2, pca5 indicates PCA with embeddings of size 2 and 5 respectively, and likewise for the other methods. For the MRMR method, mrmr30 and mrmr50 indicate selecting 30 and 50 features respectively out of the original transcripts. ga rf indicates Genetic Algorithm using Random Forest as the optimisation criteria. RF RFE refers to Random Forest Recursive Feature Elimination. While using the t-test and wilcoxontest methods, it is common to use adjusted p-value. However using adjusted p-value to select the best features was found to give poor results, therefore this graph shows the results of these methods without any adjustments for p-value. The comparison of various adjusted p-value methods is given as part of Supplementary Information. While using PCA for feature extraction, components accounting for 75% of variance was used, as this performed best among different variance thresholds. Comparison of different variance thresholds for selecting the number of components from PCA is provided in Supplementary Information.

From Figure 2, it can be seen that although L2 Regularized Logistic Regression and Random Forests have better results, the performance trend of FEMs on the datasets are approximately the same across the different classification models. PLSDA with embedding of size 5 is the best FEM for each of the different models. Comparing the datasets, it can be seen that the AUC scores for microarray-based datasets are in general on the lower side, indicating low signal-to-noise ratio in microarray datasets. An exception to this is Dataset 23, which has same order of magnitude for the number of transcripts and the number of samples as seen in Table 1. Another observation is that the TEP datasets have high performance, indicating that TEPs are effective biomarkers. This could also be attributed to the relatively large sample size of the TEP datasets used in this study.

### Statistical Analysis of Mean AUC Results

Statistical analysis of results was performed using post-hoc Nemenyi test as suggested by Demsar [45]. The Nemenyi test is a post-hoc test done after Friedman Test. Friedman Test is a non-parametric equivalent of ANOVA. For this study, the Friedman Test ranks the different feature extraction methods for each data set separately, with the best method receiving rank 1, second best rank 2 and so on. Based on these rankings, average ranking for each method is calculated. The Nemenyi test considers the performance of two methods to differ significantly if their average ranks are greater than the critical difference(CD), where 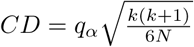 (k is the number of methods, N is the number of datasets, *q*_*α*_ based on Studentized range statistic divided by 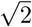). The ranking can be graphically presented using Critical Difference diagram (CD diagram). The R package scmamp [46] was used to create the CD diagrams. This approach assumes that the datasets used are independent. Some datasets used in this study originate from the same original data, nevertheless we make this dataset independence assumption.

The Critical Difference (CD) Diagram for Mean AUC for the different FEMs is shown in Figure 3. Each subfigure corresponds to a comparison of the FEMS for a specific classification model. The position of each of the methods in the CD Diagram indicates their average ranks based on performance across the 23 datasets. For the methods that are connected by horizontal lines, the difference in the average ranking is not statistically significant. For all the classification models, it can be seen that PLSDA5 is the highest ranked in the CD Diagram, however not statistically significant. But the raw ranking itself provides a means of comparison.

**Figure 3.**
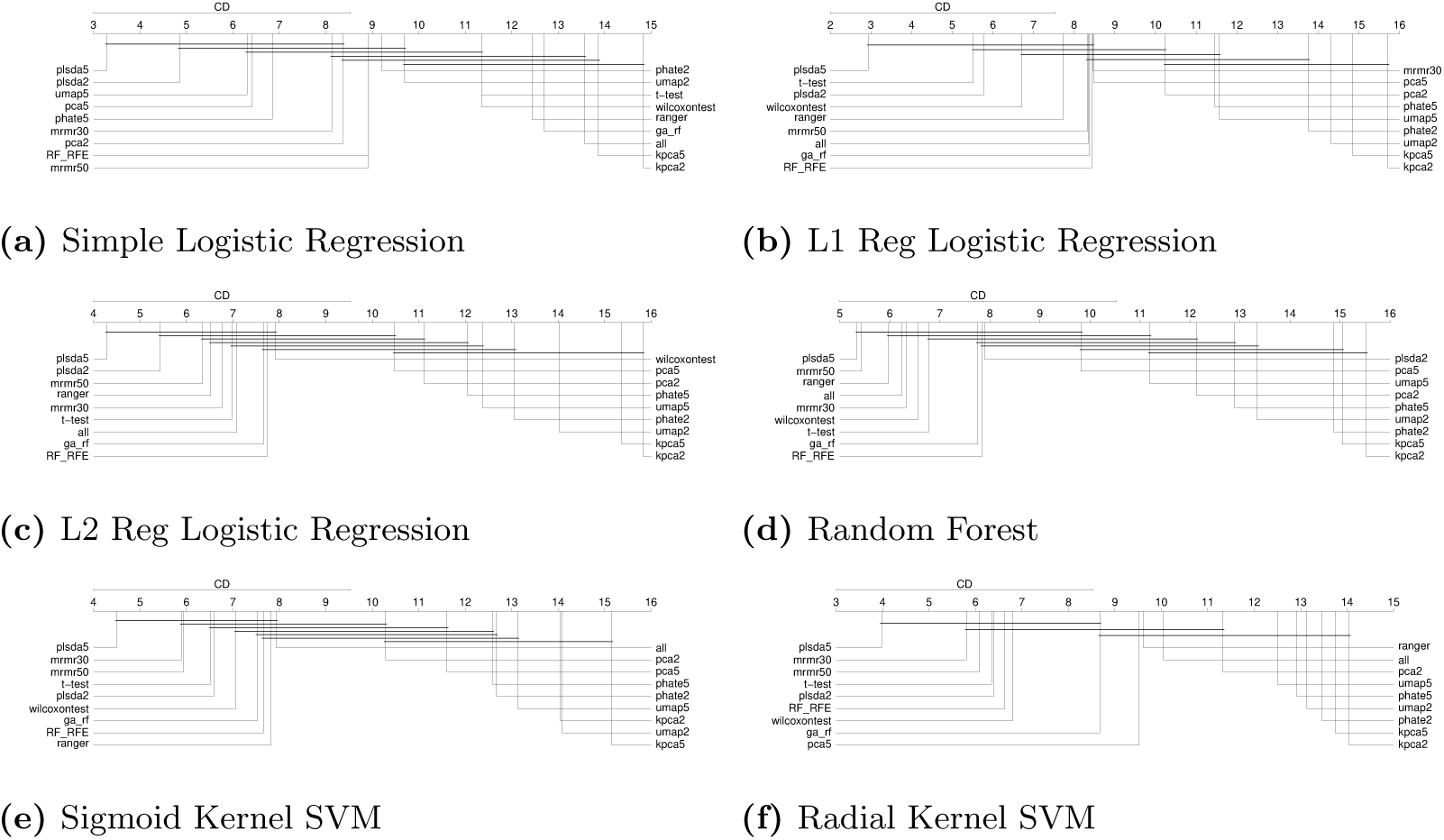
Critical Difference Diagrams comparing average ranking of feature extraction methods computed across different datasets based on their Mean AUC measure

### Analysis of number of features

The CD diagrams show that the transformation based FEMs except for PLSDA perform worse than FSMs. To verify if this is because of the small size of embeddings compared to the number of features chosen by the FSMs, an analysis on the effect of number of features was performed.

The mean number of features selected by the feature subset selection methods on the different datasets is shown in Figure 4 in log scale. The vertical black lines indicate the 95% confidence interval of the number of features. For the GSE44281 dataset, t-test and wilcoxontest fail to select any features for most of the iterations, therefore only the upper 95 CI is shown in the figure. It can be seen that the number of features selected is much higher than the embedding sizes used for the transformation methods.

**Figure 4.**
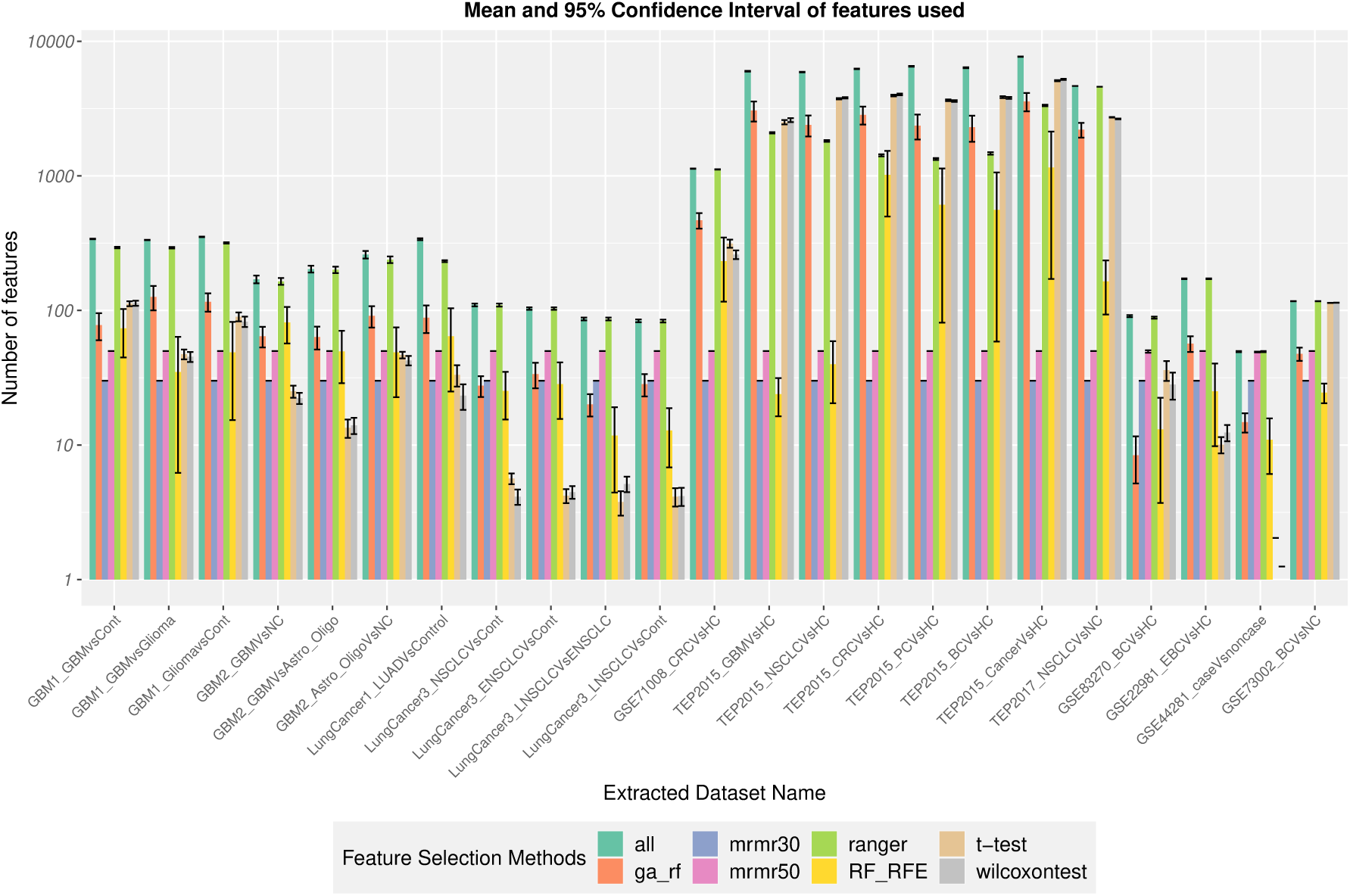
Features used in different datasets

To assess if larger embedding sizes need to be chosen, two methods of selecting larger embedding sizes were compared with the original 2 and 5 embedding sizes. 1) Embedding Size = 1 - Number of samples. 2) : Embedding Size = Embedding size required for PCA to accomodate 75% variance. 75% value is used in Method 2 because it was found that PCA with embedding size accommodating for 75% variance performed best in the datasets used.

The CD diagrams for the original embedding size and Method 1 and Method 2 are given in Figure 5. Comparing the CD diagrams in Figures 5a, 5b and 5c, it can be seen that increasing the embedding size does not improve the performance of the transformation methods, and in fact reduces it. Comparing amongst FSMs in Figure 4, it can be seen that Ranger selects the maximum number of features for non-TEP datasets, whereas for TEP datasets, the t-test, wilcoxontest, Ranger and Genetic Algorithm with Random Forest(ga rf) select large number of features, whereas Ranger selects the smallest? This implies that the transcripts in TEPs are better cancer indicators.

**Figure 5.**
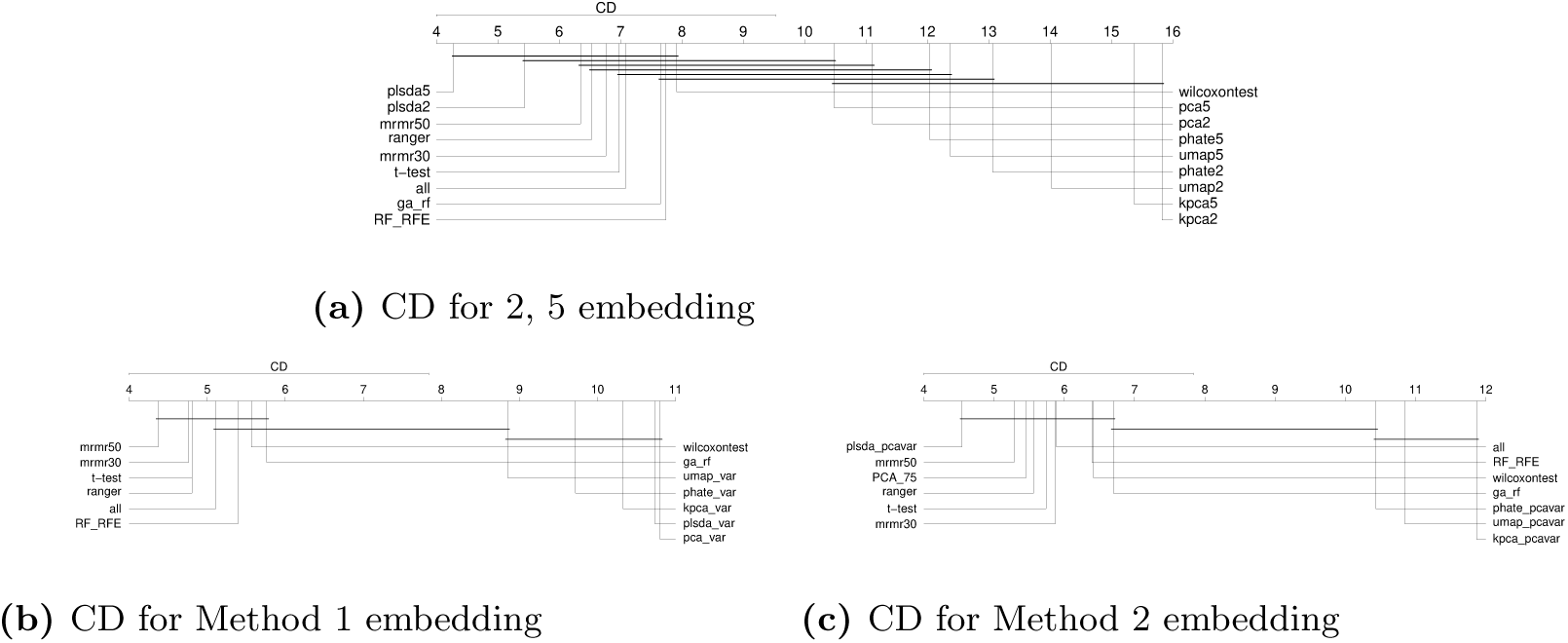
Critical Difference Diagrams using varying embedding sizes

### Effect of filtering

The filtering step during preprocessing is commonly performed in transcriptomic prediction tasks. The number of transcripts as shown in Table 1 is significantly reduced after the preprocessing filtering step. The number of transcripts after filtering can be seen in Figure 4 corresponding to using ‘all’ features. It also show that well performing Feature Subset Selection Methods have selected a large number of transcripts as good features.

Therefore, it is natural to evaluate if the methods perform better without filtering. With this goal, the best performing methods were executed without the filtering step, and their Mean AUC compared using Critical Difference diagram from Nemenyi Test as before. The resulting CD diagram for L2 Regularized Logistic Regression is shown in Figure 6. The execution of the methods without filtering is marked with suffix ‘no fil’ in the CD diagram. The figure clearly shows that each of the methods are ranked higher than the corresponding no filter version. Therefore it is clear that the filtering step does indeed aid the feature extraction process and the resultant prediction step.

**Figure 6.**
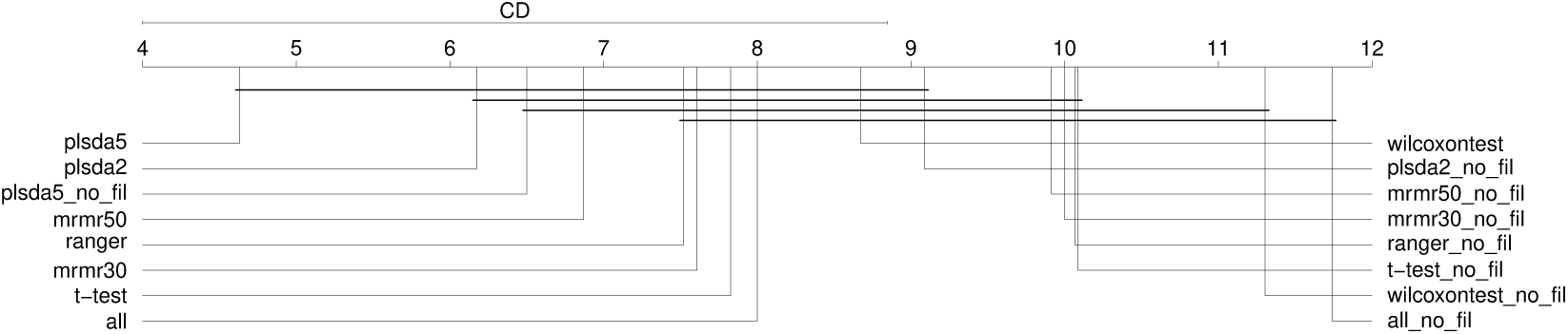
Mean AUC without filter comparison - Critical Difference Diagram

### Robustness Test

Robustness is defined as the ability for a feature extraction method to consistently provide similar performance for the classification model across multiple samples. This was evaluated by verifying if 1) FEMs select similar features 2) FEMs provide comparable AUC

#### FEMs select similar features

Testing for similar features selected can be performed only on Feature Subset Selection Methods. Average of pairwise Jaccard Index (JI) over 30 iterations of train-test split was used as the evaluation metric.

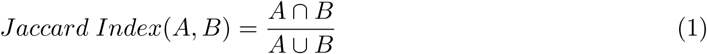

where:

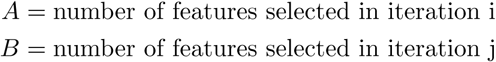

Average pairwise JI for each of the Feature Subset Selection Methods on all the datasets is shown in Figure 7. It can be seen that among the Feature Subset Selection Methods (i.e. excluding selecting all features), Ranger has the best pairwise Jaccard Index and hence is the most robust method.

**Figure 7.**
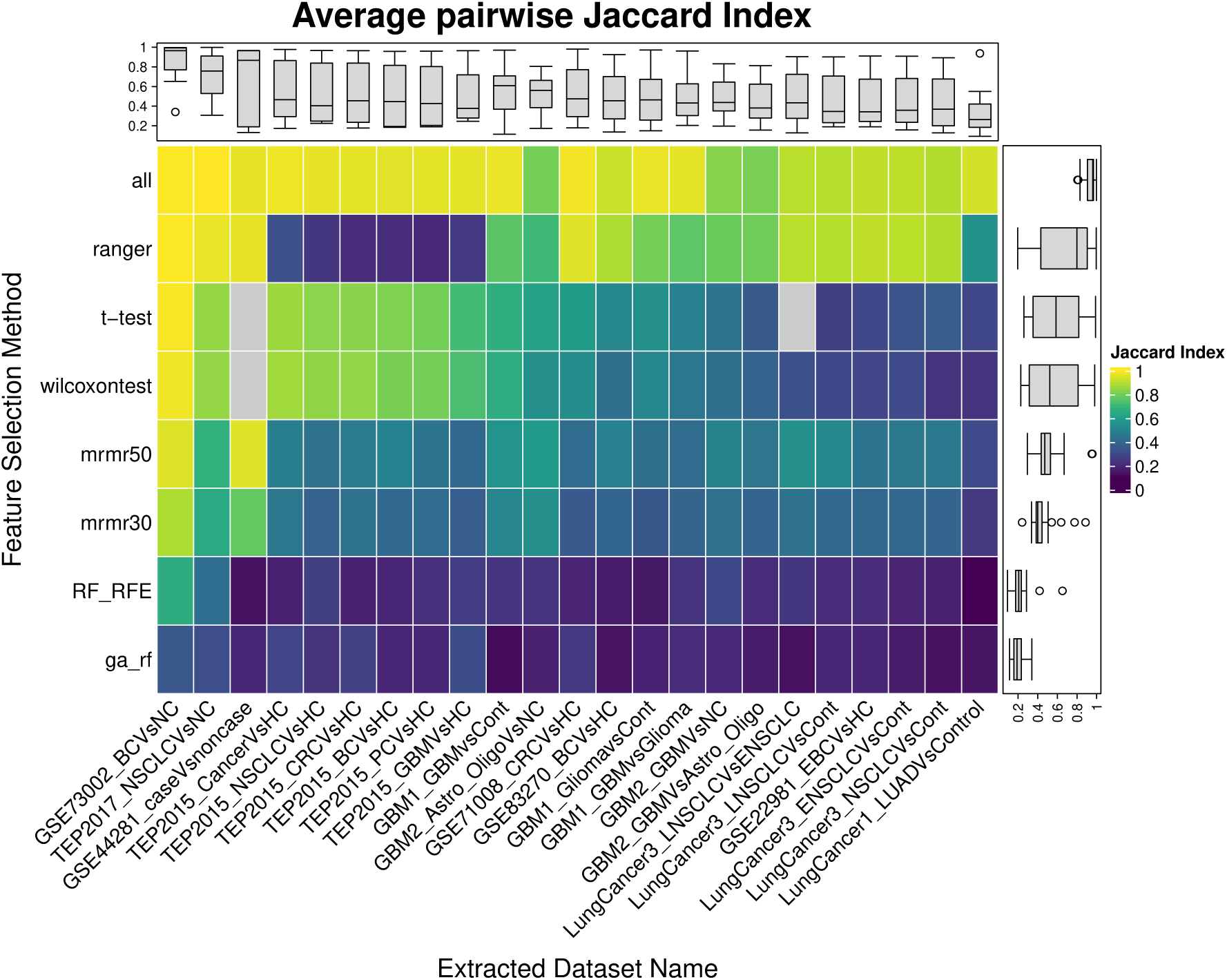
Average pairwise Jaccard Index

Note that the JI value for selecting all features is not perfectly 1 because the preprocessing filter step provides different sets of transcripts for different train-test splits.

#### FEMs provide comparable AUC

To evaluate the robustness, taking into account the transformation based feature extraction methods, the difference in 95 percent Confidence Interval (95 CI) of the AUC is compared in Figure 9. This figure shows the AUC values using L2 regularized logistic regression model. In this figure, the vertical black lines in each of the bars corresponding to the FSMs connect the lower and upper 95 CI. Hence the length of the vertical black lines indicate the difference in 95 CI. This difference in 95 CI is then ranked using Nemenyi test in the CD Diagram in Figure 8. The method with the worst ranking will be the best in terms of lowest 95 CI. In Figure 8, Ranger performs third best on this measure of robustness. The best performing feature extraction method in terms of AUC is PLS-DA, which is not far behind and comes seventh in robustness ranking.

**Figure 8.**
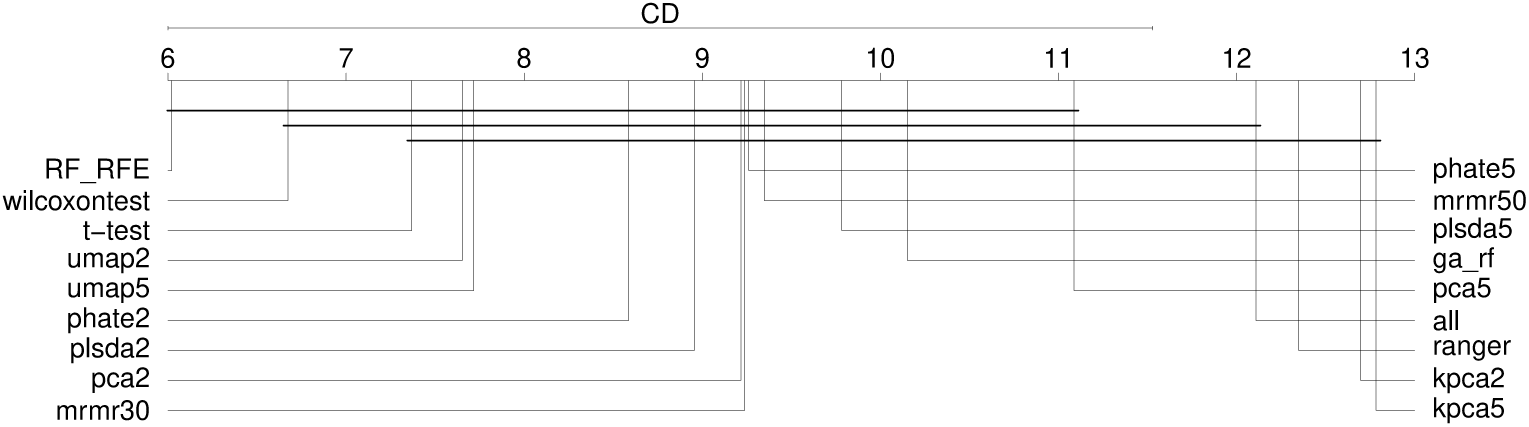
Difference in 95 CI - Critical Difference Diagram

**Figure 9.**
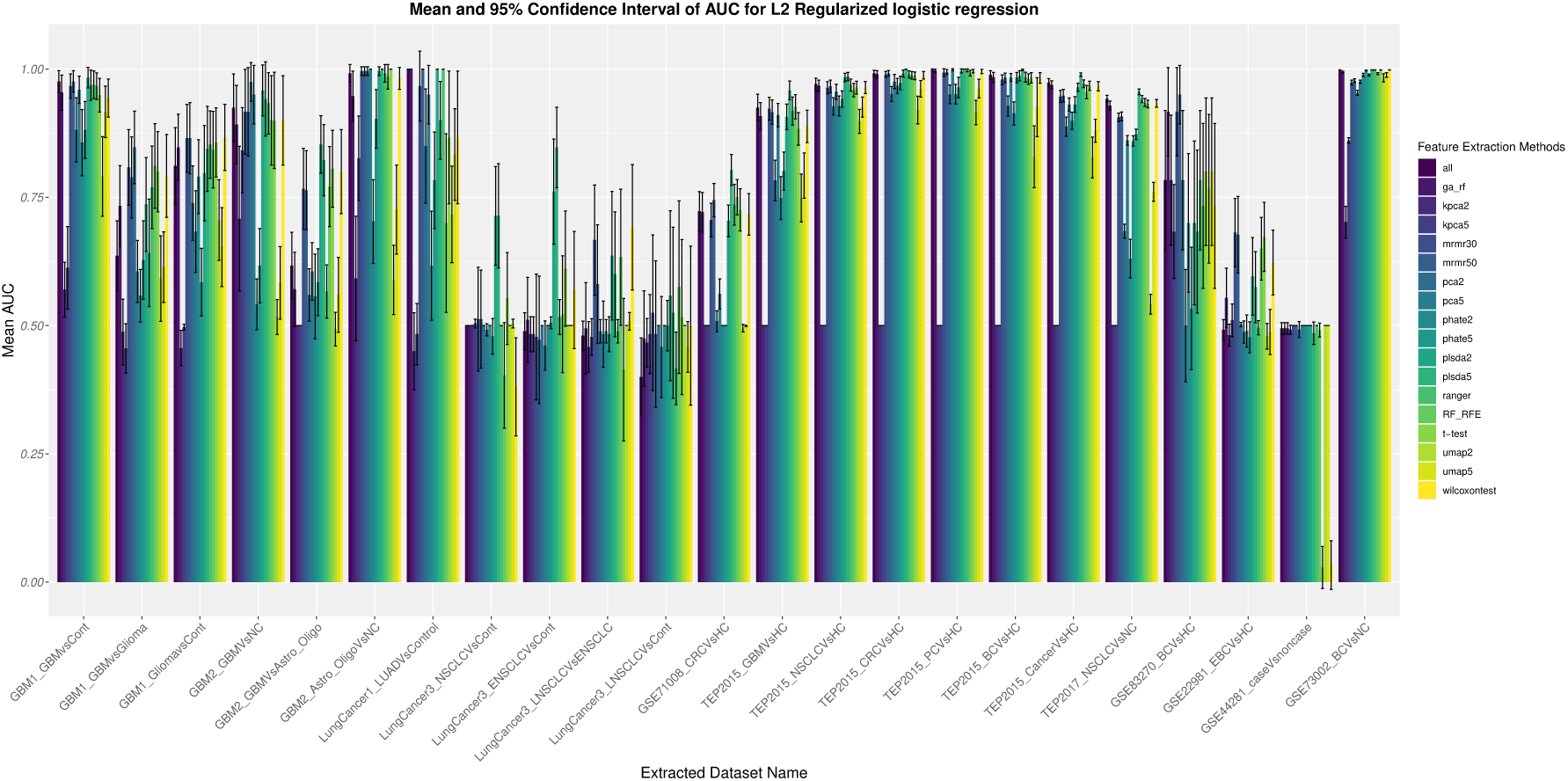
Difference in 95 CI of AUC

## Conclusion

In this study, a generic pipeline was created to perform preprocessing, feature extraction and classification steps for cancer prediction from liquid-biopsy based transcriptomic data. Through the benchmarking performed in this work, PLS-DA with embedding size of 5 (PLSDA5) was found to be the best performing feature extraction method across all models in terms of Mean AUC. PLSDA5 is well performing in terms of robustness, computed using variability in performance, across different samples as well. Ranger, which is found to select a large number of features, performs the best in terms of robustness, computed using Jaccard Index of selected features. It has also been established that the preprocessing step of filtering aids in the feature extraction process and the resultant prediction. The generic pipeline created in this work and the associated R package will aid in effortlessly extending the study to other omics datasets.

The findings from this study need to be further validated by assessing the performance on external datasets. Other tests of robustness such as robustness to random elimination of features and robustness to addition of noise can be performed. Currently Neural Networks based approaches have not been attempted, primarily because of lack of sufficient samples. Once sufficient datasets / samples are available, Deep Neural Network architectures can be explored. Further investigation is also required to ascertain why Ranger performs well, even though it selects almost all features.

## Data and Code availability

All datasets used in this study are publicly available with GEO accession numbers provided in Table 1 The pipeline is fully-implemented in R and is generalisable to other omics databases. The corresponding R package is freely available for non-commercial uses in GitHub (first release DOI: https://doi.org/10.5281/zenodo.6300985). Future updates of the R package will be available on GitHub: https://github.com/abhivij/bloodbased-pancancer-diagnosis

## Author contributions statement

FV conceived and supervised the study, envisaged the overarching goal, directed the methodology, and critically reviewed and revised the manuscript. AS provided guidance to AV on methods, experimental design and results analysis. AV wrote the code for the pipeline and created the R package, performed the analysis, wrote the initial draft and the revised version of the manuscript. SF helped AV with data collection, understanding the biology and contributed to the manuscript writing. All authors reviewed the manuscript and approved the final version.

## Notes

### Competing Interest Statement

The authors have declared no competing interest.

