## Supplementary Information for "Blood-based transcriptomic signature panel identification for cancer diagnosis: Benchmarking of feature extraction methods"

### Supplementary Figures

**Supplementary Figure 1.** Adjusted p-value comparison.

**Supplementary Figure 2.** Variance threshold comparison for PCA.

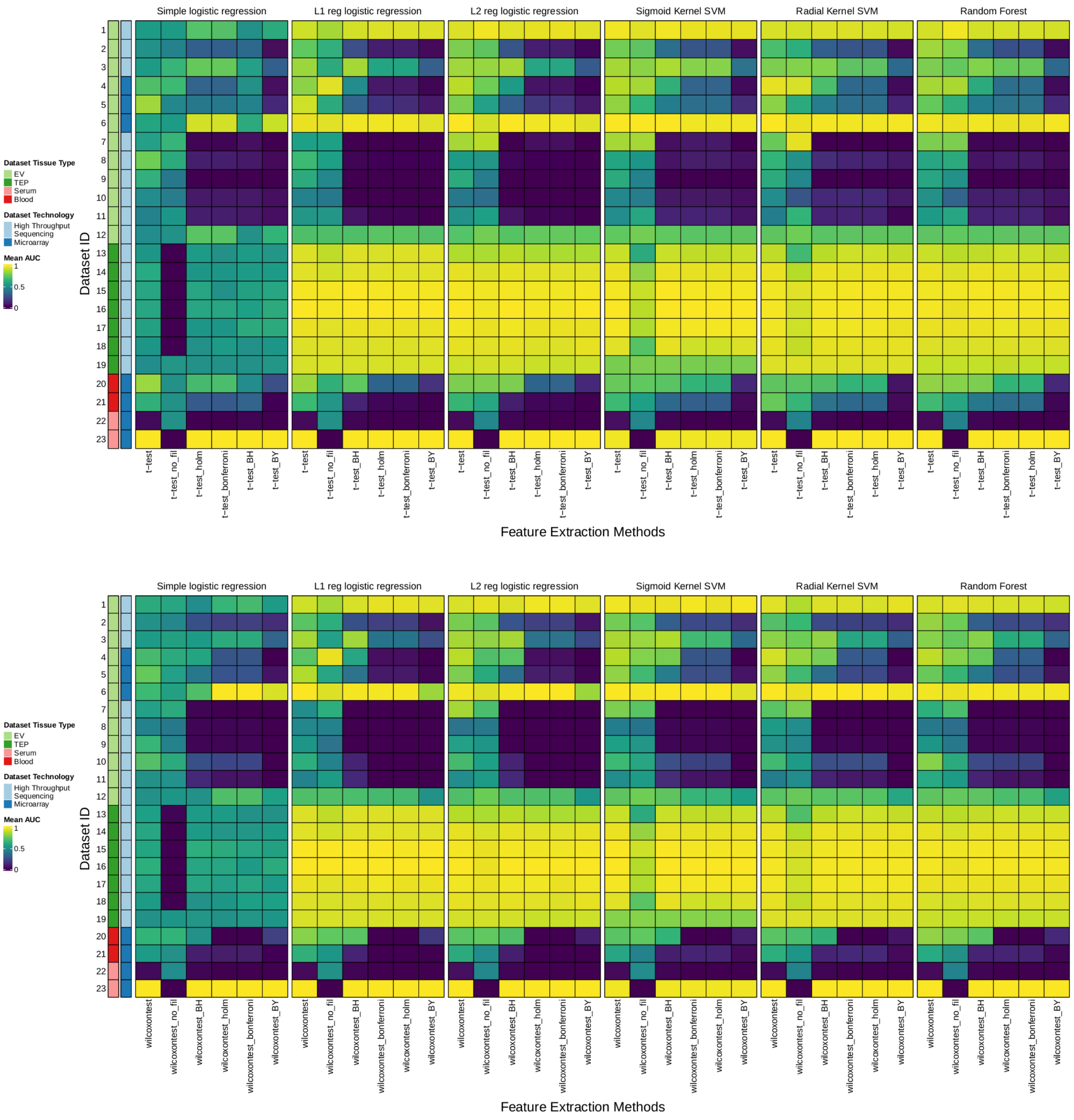


**Supplementary Figure 1.** The above two figures show various adjustment methods applied on the obtained p-value from for t-test and Wilcoxon test. Also shown are the methods without the filtering step denoted by the no_fil suffix. It can be seen that the original methods (i.e., t-test and Wilcoxon test) with filtering perform the best.

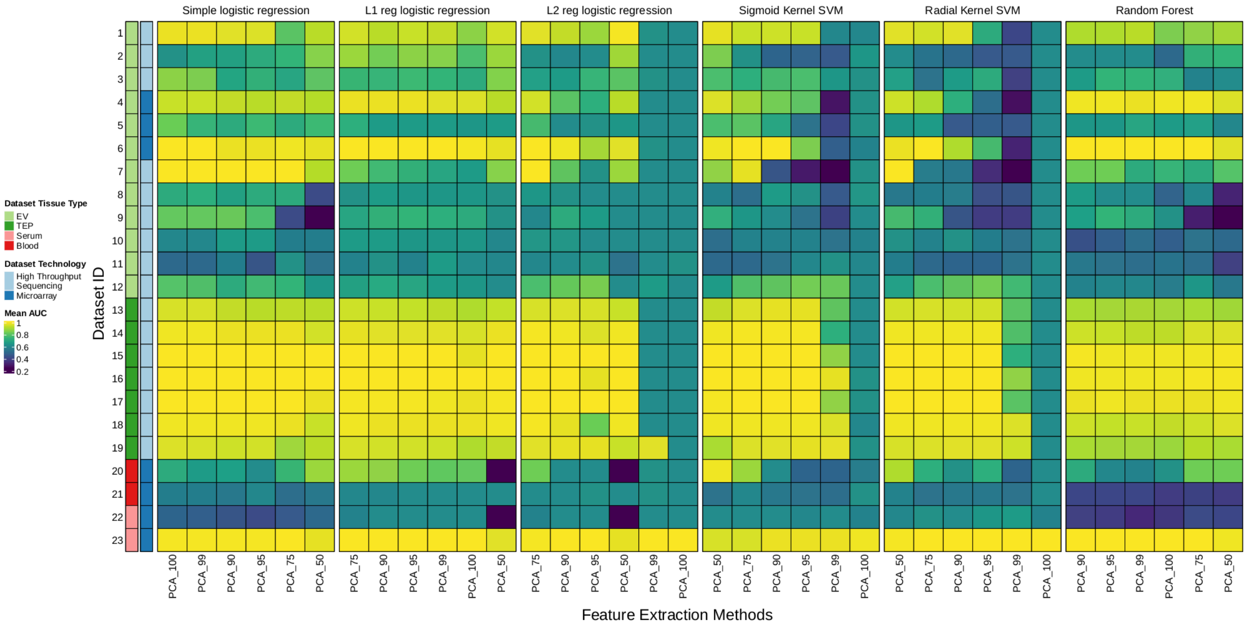


**Supplementary Figure 2.** From the above figure it can be seen that among the different variance thresholds, PCA with the number of components accommodating for 75 % variance performs best in a maximum number of models compared to others.
